# Dynamic transcriptional heterogeneity in pituitary corticotrophs

**DOI:** 10.1101/2025.04.04.645979

**Authors:** Zhengyang Yuan, Linda Laiho, Elsa Smith, Craig McDougall, Peter J Duncan, Paul Le Tissier, Michael J Shipston, Nicola Romanò

## Abstract

A large body of evidence has shown that corticotrophs, the anterior pituitary cells central to the generation of hormonal stress responses, exhibit heterogeneous functional behavior, suggesting the presence of functional sub-populations of corticotrophs. We investigated whether this was the case at the transcriptomic level by conducting a comprehensive analysis of scRNA-seq datasets from rodent pituitary cells. We envisaged two alternative scenarios, one where robust subtypes of corticotrophs exist, and the other where these subpopulations were only transient states, possibly transitioning into one another. Our findings suggest that corticotrophs transition between multiple transcriptional states rather than existing as rigidly defined subpopulations. We employed marker gene-based comparisons and whole transcriptome label transfer approaches to analyze transcriptional signatures across datasets. Marker-based clustering revealed strikingly low similarity in the identified subpopulations across datasets. This analysis evidenced the presence of transcriptional states with different functional relevance, related to different stages of hormonal signalling. Similarly, the label transfer approach, which considers non-linear interactions across the entire transcriptome showed that transcriptional states could be detected across independent datasets. This classification relied on broader gene expression patterns rather than conventional marker genes, reinforcing the notion of continuous rather than discrete cell states. Furthermore, trajectory analysis by RNA velocity indicated dynamic transitions between transcriptional states, suggesting the presence of transcriptional mechanisms facilitating rapid recruitment of corticotrophs in response to physiological demands.

Our findings align with evidence from other endocrine cell types, such as lactotrophs and pancreatic β-cells, where hormone secretion is linked to fluctuating transcriptional activity. The observed transitions in corticotroph states suggest a mechanism allowing flexible hormonal responses to unpredictable and time-varying stressful events. Additionally, this study highlights the challenges associated with scRNA-seq methodologies, including data sparsity, batch effects, and pseudoreplication, underscoring the need for rigorous experimental design and reproducibility in single-cell transcriptomics research. These insights contribute to a broader understanding of pituitary cell plasticity and endocrine adaptation mechanisms.

## Introduction

Heterogeneous cell behaviour is an intrinsic property of cellular systems (Huang, 2009; Raj and van Oudenaarden, 2008). While heterogeneity is driven by the stochastic nature of transcription and other cellular events, it is not merely a source of noise, but it fulfils critical biological functions. The role of heterogeneity is particularly evident in endocrine tissues, where diverse cell populations must coordinate their activity to maintain physiological homeostasis. The pituitary gland is a prime example of such an organ, with distinct yet heterogeneous cell populations. Complex interaction dynamics between the five hormone-producing cell types and non-endocrine cells in the gland, in response to hypothalamic and peripheral inputs, control key physiological processes, such as growth, reproduction and stress (Le Tissier et al., 2017). These cells must respond to changes in the internal and external environment to maintain body homeostasis through precisely controlled production of hormones; thus, this requires the coordinated action of pituitary cell populations. However, heterogeneity has been shown as a prominent feature of neuroendocrine systems, raising the apparent paradox of how heterogeneous single cells come together to generate coordinated population outputs proportionate to environmental challenges. The variety of possible amplitude, duration and type of challenges, which are not always predictable, and the dependence of cell responses on physiological status make cell coordination in the gland complex (Le Tissier et al., 2022).

This study focuses on anterior pituitary corticotrophs, which secrete adrenocorticotropic hormone (ACTH) in response to stress. Heterogeneity in this cell population was noted in early studies where differences in corticotroph morphology were revealed by electron microscopy (Yoshimura and Nogami, 1981). Single-cell secretion studies further evidenced that single corticotroph cells exhibit variability in the amount of ACTH secreted following a stimulus; increasing the strength of the stimulus results in the recruitment of a higher proportion of cells in the population (Lugo and Pintar, 1996; Canny et al., 1992; Childs and Burke, 1987). Similar behaviour has been reported when analysing firing behaviour (Liang et al., 2011), intracellular calcium changes in response to physiological stimuli (Romanò et al., 2017; Corcuff et al., 1993), or activation of intracellular pathways (Takigami et al., 2008). This points towards the idea that sub-populations of corticotrophs could exist, possibly with different functional roles; for example, certain subsets of cells might only be recruited when stronger or longer stressors are sensed, thus increasing the dynamic range of the system.

Recent advances in sequencing technology have allowed an increasingly detailed and fine-grained view of the transcriptional activity of single cells. Single-cell RNAseq (scRNAseq) experiments have confirmed that the phenotypic heterogeneity of pituitary tissue described above is also reflected at the transcriptional level. Several studies have brought forward the idea of transcriptional subpopulations of hormone-producing cells (Sheridan et al., 2024; Cheung et al., 2018, 2023; Zhang et al., 2022; Ruf-Zamojski et al., 2021), including two subpopulations of human corticotrophs (Zhang et al., 2020). While evidence of transcriptional subpopulations of these cells is compelling, it is often limited to single studies and a comprehensive analysis of corticotroph transcriptome and its heterogeneity at the single-cell level is currently lacking.

We thus hypothesised that sub-populations of corticotrophs exist and envisaged two alternative scenarios, one where robust, stable sub-types of corticotrophs exist, and the other where these subpopulations were only transient states, possibly transitioning into one another. To test our hypothesis, we re-analysed publicly available scRNAseq datasets of pituitary gland tissue from rodents to explore corticotroph heterogeneity and the presence of these subpopulations. If there were robust sub-types of corticotrophs, it should be possible to detect them in similar proportions in any of the pituitary scRNAseq datasets analysed, and possibly even between different species. On the other hand, if these were dynamic cell states rather than stable sub-populations, we would expect to detect them in different proportions in different studies and possibly in only some of them.

This type of comprehensive analysis is challenging due to the variety of approaches used by different studies and the problem of objectively defining what a cell type or subtype is (Fleck et al., 2023; Zeng, 2022), especially in a small population of cells such as corticotrophs.

To overcome some of these issues, we used a variety of bioinformatics approaches to identify subpopulations in publicly available pituitary datasets, and match possible subpopulations or cell states between different studies. From our results, we conclude that corticotrophs can exist in several dynamic transcriptional states, which can transition into one another. This plasticty could be a mechanism to rapidly adapting to unpredictable changes in the external environment and enhance the population response to different stressors.

## Methods

### scRNAseq datasets

We performed a comprehensive literature search for all articles in which scRNAseq was performed on the pituitary gland as of January 2023, restricting our analysis to rodent datasets and to non-stressed conditions only to have as comparable a set of data as possible. We only considered mouse and rat data, from both males and females; we decided to exclude human datasets since the vast majority analyse only tumoural tissue, and avian or fish datasets as they are scarce.

### Alignment to the genome and quality control

FASTQ files for the control animals in each of the datasets were downloaded from public repositories (Supplementary Table 1), and reads were aligned to the mouse mm10 GENCODE vM23/Ensembl 98) or the rat mRatBN7 genome using CellRanger v6.1.2. Data from males and females were analysed separately.

The filtered outputs from CellRanger were imported into R 4.2.2 using the Seurat 4.3.0 package (Hao et al., 2021). After inspection of quality control (QC) parameters, including the number of UMI/cell, genes/cell and percentage of mitochondrial genes (identified as genes with MGI identifier starting with mt-), filtering was performed to remove multiplets and low-quality cells. See the Results section for a detailed description of the QC process.

Data was scaled using regularised negative binomial regression (SCT, (Hafemeister and Satija, 2019)), independently in each dataset, and dimensionality reduction was performed through Principal Component Analysis (PCA), keeping the first 50 principal components; cells were projected onto a 2D space using Uniform Manifold Approximation and Projection (UMAP) for visualisation purposes.

### Cell typing

We identified corticotrophs as cells expressing high levels of the *Pomc* gene, coding for the hormone ACTH but not expressing *Pcsk2* (coding for the proprotein convertase type 2, which converts ACTH to α-MSH in melanotrophs) or the key regulator of melanotroph identity *Pax7*. To determine the threshold used to distinguish cells expressing each of these genes, we used Otsu’s method (Otsu, 1979) on the distribution of counts for the two genes. Given that there are two cell populations (e.g. *Pomc* positive and negative), Otsu’s method finds the optimal threshold *t* that minimises the intra-class variance defined as:

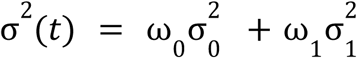

Where σ^2^*_i_* is the variance of class *i*, weighed by a factor ω*_i_*.

Corticotrophs were isolated from the data and were sub-clustered using Louvain’s algorithm, using k=20 for the k-nearest neighbours algorithm, using the Seurat R package. The results of this procedure are dependent on the clustering resolution, with lower resolution values generating fewer, more heterogeneous clusters. The optimal clustering resolution was chosen using a bootstrapping procedure. We calculated the average within-cluster sum of squared errors (WSS) at different resolutions ranging from 0.1 to 1.5 in steps of 0.1 for each dataset. The WSS was calculated over 100 rounds of clustering performed on a random 60% bootstrap sample of the population. We then used the “elbow method” to choose the resolution at which the WSS stopped decreasing (showing an “elbow” in the WSS plot). The results of this procedure are shown in Supplementary Figure 1.

### Marker genes and pathway analysis

Marker genes for each cluster were calculated in Seurat using a Wilcoxon sum rank test, filtering only for genes showing at least 0.25 ln-fold change increase in at least 20% of cells. Pathway analysis was performed using ShinyGO 0.82, using the full list of genes common to all datasets as background and querying for FDR-corrected enrichment of GO:Biological Process, GO:Molecular Function and KEGG pathways (Ge et al., 2020).

Module scores for enriched pathways were calculated using the *AddModuleScore* Seurat function (Tirosh et al., 2016); gene lists for each pathway were obtained using the *KEGGREST* 1.46 (Tenenbaum, 2024) and *biomaRt* 2.62.1 (Durinck et al., 2009) R packages; in the case of early response genes, a list was obtained from (Tullai et al., 2007). Pairwise comparisons between communities were performed using a bootstrap resampling approach (n = 1000 iterations). In each iteration, stratified resampling with replacement was conducted at the dataset level to account for nested data structure, and median pathway scores were computed for each community. The p-value for each comparison was estimated as twice the smaller proportion of bootstrap samples where the observed difference in median pathway scores was either greater or less than zero, to approximate a two-tailed test.

### Correlation analysis

Spearman’s correlation of the level of expression of markers of each dataset was calculated (see Results) and the level of correlation was compared using a mixed effect model, calculated using the *lme* function of the *nlme* 3.1 R package (Pinheiro and Bates, 2000); fixed effects included sex of the animals, a variable indicating the group, with levels “Within clusters”, “Between clusters”, “Random shuffle” and “Random shuffle within”, and a variable indicating whether the comparison was calculated on the same dataset or not. Multiple comparisons were calculated using the *emmeans* 1.10.7 R package.

### Label transfer

Expression matrices (SCT-transformed values, filtered only to include the 11103 genes detected in all datasets) and cluster labels were exported from Seurat and loaded into Python 3.11.4. Data were split into a training and test set, with a 90%/10% ratio; the test set was held out and never used during training or hyperparameter optimisation. During training, the training set was further split into a training (80%) and validation (20%) set. The training sets were used to train a series of multi-layer perceptrons, one for each dataset, using the Keras framework (version 3.3.3). The general structure of these classifiers was an input layer, followed by a batch normalisation layer, one hidden layer with ReLU activation and L1 regularisation, followed by a softmax output layer. The macro-F1 score, defined as the harmonic mean of precision and recall for each class, was used as an evaluation metric, together with the area under the curve of the receiver-operating characteristic curve (ROC AUC).

Hyperparameters were initially tuned using an exhaustive grid-search strategy, iterating over possible values of two hyperparameters at a time; we started by optimizing the number of nodes in the hidden layer and the learning rate, then the other hyperparameters. Initial tests with multiple hidden layers showed that adding depth led to overfitting and decreasing F1 scores, possibly because of the small number of cells in these datasets. Five-fold cross-validation was used during hyperparameter tuning to avoid overfitting and improve model generalisation. Upon visual inspection of the training curves, further adjustment of the hyperparameters was performed manually to minimise overfitting. This included adding batch normalisation, dropout, and L1 normalisation in the hidden layer. The final hyperparameters chosen for each dataset are shown in Supplementary Table 4. After hyperparameter tuning, the models were retrained on the whole training dataset to maximise the amount of data and finally evaluated on the held-out test set. The final F1 scores and ROC AUC evaluated on the held-out test set are reported in Supplementary Table 5.

Each trained model was used to predict all other datasets. Cells for which the predicted probability of the top class was less than 0.7 were considered unassigned.

Feature importance for the prediction was determined using Deep Shapley additive explanations (DeepSHAP) (Lundberg and Lee, 2017) as implemented in the *shap* Python package.

### Cluster communities

Graphs of the cluster relationships between datasets were generated using iGraph 0.10 in R. Node communities were defined using the Walktrap algorithm (Pons and Latapy, 2005); an identifier of the community was saved in the metadata for further analysis. These can be accessed in the provided .rds files (see Code and data availability below) through the *marker_community_<percent>* or *nn_community_* metadata.

### RNA velocity analysis

RNA velocity was performed using scVelo 0.2.5 (Bergen et al., 2020). The CellRanger output was filtered only to include reads from corticotrophs based on the cell barcodes obtained during the preprocessing steps described above. Velocyto (La Manno et al., 2018) was used to generate loom files and velocity data was then visualised on UMAP embeddings, annotated with the communities found from the previous analyses. A directed graph was then constructed to show the transitions between different communities. To do so, the average velocity vector was calculated for each community (calculated using either of the methods described above) and a transition matrix was built with the cosine similarity between the average velocity of each community 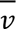 and the vector *d* between the median centroid of that community and that of all other communities, scaled by the magnitude of the velocity vector, calculated as:

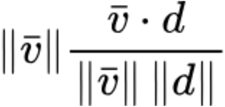

This matrix was thresholded for values of similarity greater than 0.75 (corresponding to an angle of ∼40°) to determine communities transitioning toward each other. The normalised transition matrices, weighted by the number of cells in each dataset, were then used to produce a directed graph using iGraph.

### Code and data availability

All the code for the analyses performed in this manuscript is available on GitHub at https://github.com/nicolaromano/corticotroph-heterogeneity. The code has been annotated to make it as easy as possible to reproduce the analyses in this article or generalise them to other datasets in a stepwise manner. Trained models for label transfer, expression matrices and cell labels can also be found in the same repository, along with files for each dataset containing the Seurat object with complete metadata. We also provide R (*renv*) and Python (*venv*) environment files to allow users to recreate our analysis environment with the exact versions of software, libraries, and dependencies used in this manuscript.

## Results

### Data preprocessing

We identified seven datasets containing scRNAseq data from mouse pituitary and two from rat pituitary (Table 1) that we used for our study. The full metadata, including links to the publicly available raw data, are listed in Supplementary Table 1. Only control animals have been considered in this study, and owing to the known sex differences in corticotrophs (Duncan et al., 2023; Heck and Handa, 2019; McCarthy et al., 2012), data from male and female animals were analysed separately. Due to the low number of rat dataset we decided against analysing those separately.

**Table 1.**
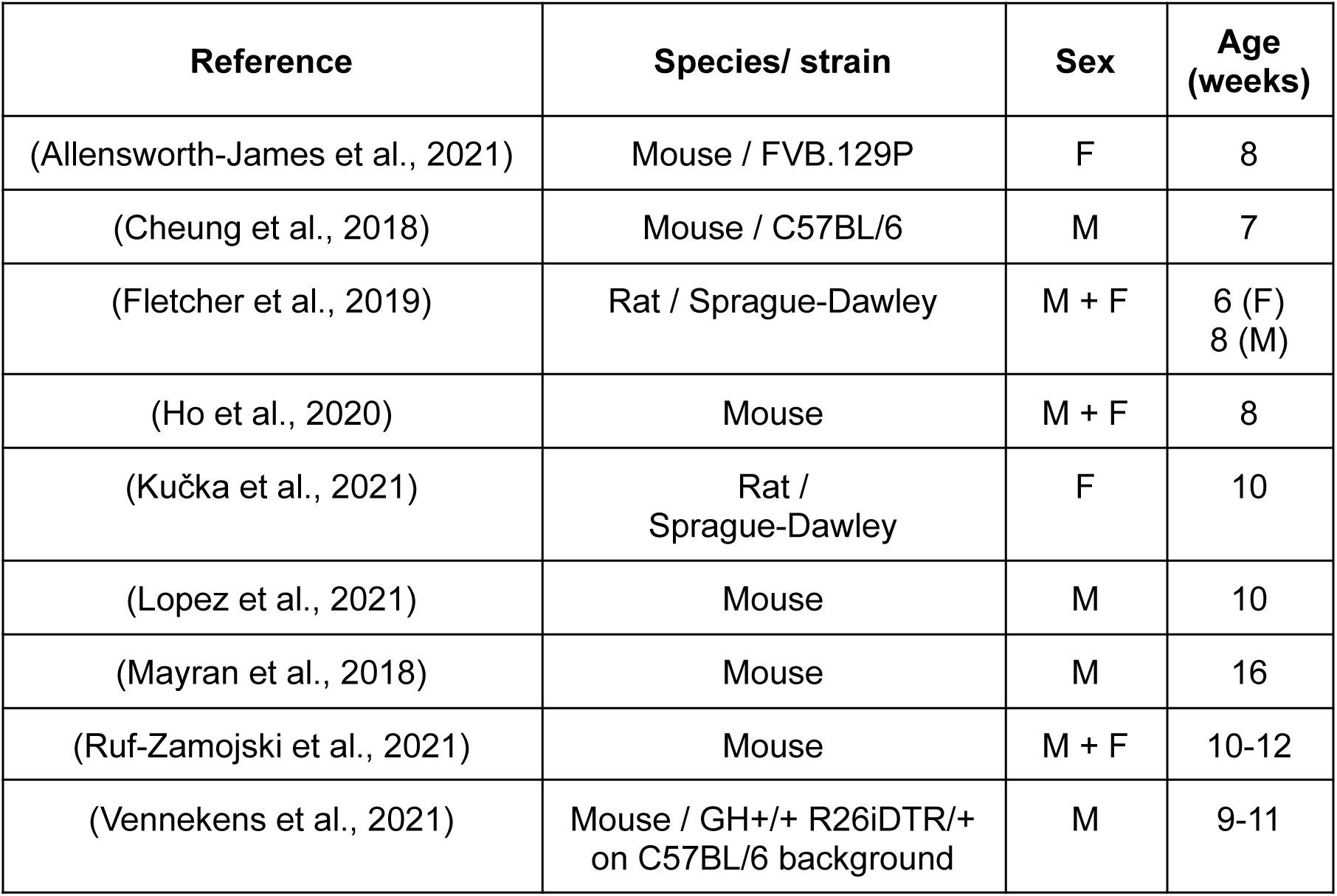
List of the datasets used in this analysis, including strain, sex and age of the animals used. Only data from control animals were used.

| Reference | Species/ strain | Sex | Age (weeks) |
| --- | --- | --- | --- |
| (Allensworth-James et al., 2021) | Mouse / FVB.129P | F | 8 |
| (Cheung et al., 2018) | Mouse / C57BL/6 | M | 7 |
| (Fletcher et al., 2019) | Rat / Sprague-Dawley | M + F | 6 (F)<br>8 (M) |
| (Ho et al., 2020) | Mouse | M + F | 8 |
| (Kučka et al., 2021) | Rat /<br>Sprague-Dawley | F | 10 |
| (Lopez et al., 2021) | Mouse | M | 10 |
| (Mayran et al., 2018) | Mouse | M | 16 |
| (Ruf-Zamojski et al., 2021) | Mouse | M + F | 10-12 |
| (Vennekens et al., 2021) | Mouse / GH+/+ R26iDTR/+<br>on C57BL/6 background | M | 9-11 |

After aligning the data to the genome, we performed quality control (QC) to eliminate low-quality cells. QC criteria used in the different studies were diverse (as listed in Supplementary Table 2); to ensure consistency in the processing of the data and allow comparisons between datasets, we decided to use a unified set of criteria. Despite our best efforts either using the criteria from the original papers or different combinations of QC parameters, we could not find a set of QC parameters that consistently recapitulated the number of cells reported in the publications by each of the studies; thus, we decided on a set of criteria which filtered what we would consider low-quality cells (Figures 1A, B). We removed cells with more than 70% mitochondrial gene content, those where the number of detected genes was lower than the 25th percentile minus three standard deviations of the distribution in the population, or 500 genes, whichever was higher. Similarly, we removed cells with a number of genes exceeding the 75th percentile plus three standard deviations. Finally, cells with a number of UMI lower than the 25th percentile minus three standard deviations or 300 UMI, whichever was higher. We only considered genes detected in all the studies, resulting in 11103 features in each dataset. This resulted in a number of cells in each dataset that, overall, was not statistically different to those reported in the original publications (p=0.127, linear model, Figure 1A, B) in both sexes (% not dependent on Sex, p=0.494, linear model); however, while there was no overall statistical difference, for some of the studies the number of cells we obtained deviated notably from what was reported in the corresponding publication, highlighting how the choice of pre-processing could strongly affect analysis outcomes. Figure 1C shows the three QC features in all datasets after filtering. Data from (Allensworth-James et al., 2021) and (Ho et al., 2020) showed a low number of counts/cells and genes/cells, the latter also showing a lower number of cells.

**Figure 1.**
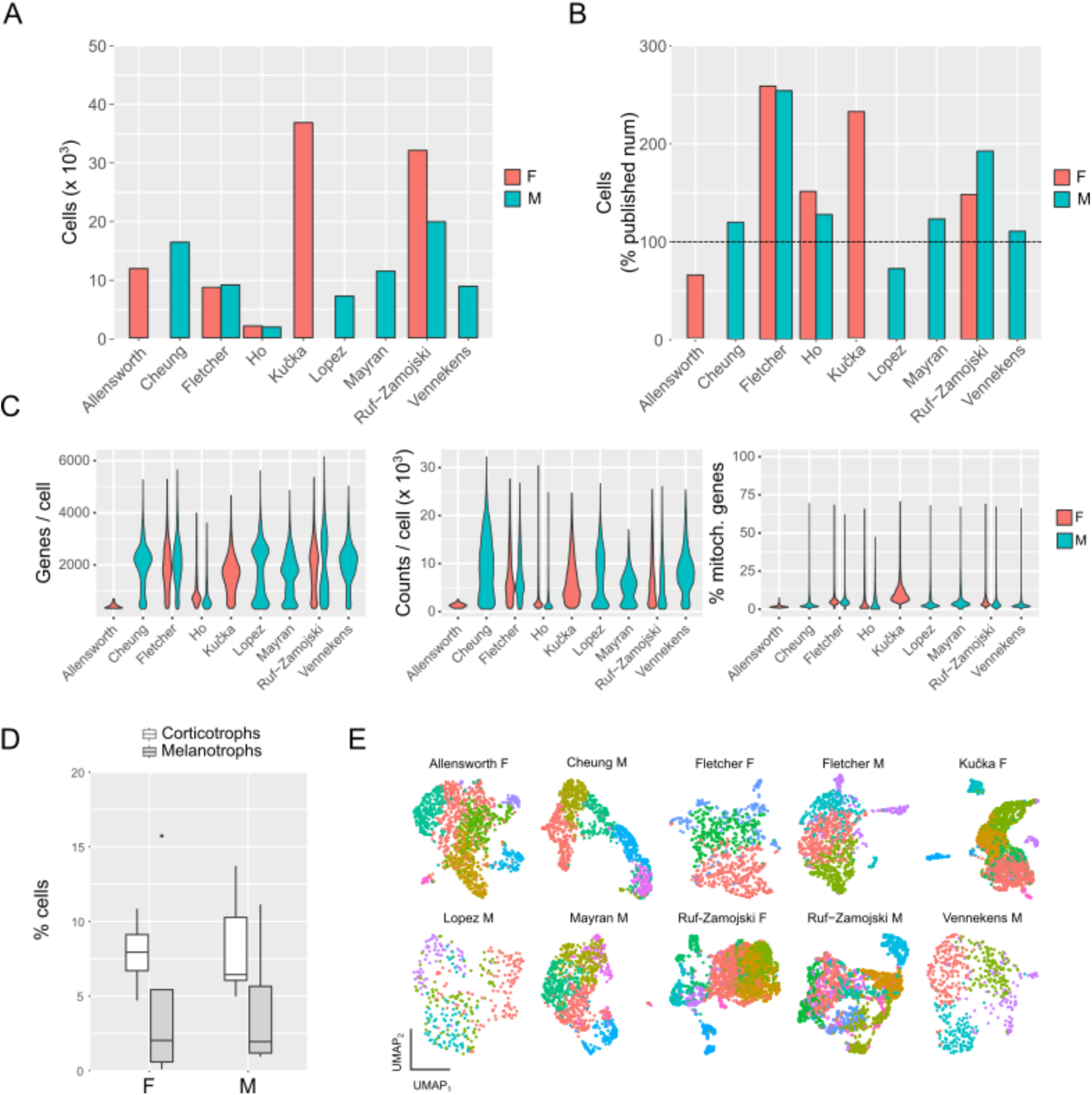
Preprocessing of the datasets analysed in this study. A) Absolute number of cells obtained following our QC filtering and B) percentage of this number with respect to what is reported in the publications (overall, not statistically different from 100%, p=0.127, linear model). C) Distribution of the 3 QC parameters used (genes/cell, counts/cell and percentage of mitochondrial genes), after filtering. D) Percentage of the total cells represented by corticotrophs (white bars) or melanotrophs (grey bars) in male vs female samples. E) Louvain cluster analysis of all datasets shown on the corresponding UMAP embeddings. Each dataset was analysed independently, so colours are not matched between different studies.

### Annotation of corticotrophs across datasets

To define corticotrophs, we selected cells with high expression of the *Pomc* gene, which codes for the ACTH precursor pre-opiomelanocortin (POMC), but that did not express *Pcsk2*, coding for the proprotein convertase 2 enzyme or *Pax7*, genes expressed by melanotrophs of the pars intermedia but not by corticotrophs (Budry et al., 2012).

Expression of these genes was binarised using Otsu’s thresholding, and cells positive for *Pomc* but negative for *Pcsk2/Pax7* were isolated for further processing (Supplementary Figure 1A; see Methods for details). In most datasets, corticotrophs and melanotrophs clustered separately on UMAP embeddings, suggesting that the procedure successfully identified these cells (Supplementary Figure 1B). Corticotrophs represented between 5 and 10% of the total cell population, in agreement with what is reported in the literature (Surks and Defesi, 1977); there was no sex difference in the percentage of corticotrophs (t_7.47_=0.814, p=0.441, two-sample Welch t-test, Figure 1D). Given the low coverage in the dataset from (Ho et al., 2020) and that the number of corticotrophs detected was low compared to the other datasets (Figures 1A and 1C), we did not analyse those data further. Figure 1E shows the UMAP projection of the corticotroph data in all datasets after sub-clustering each dataset independently using Louvain’s method by choosing the resolution that minimized the within-cluster sum of squared errors while balancing cluster granularity, as determined by the elbow method (see Methods and Supplementary Figure 1C for how clustering resolution was chosen).

### Low similarity in marker genes between subclusters indicates transient cell states

Analysis of marker genes is a popular method to define cell types in scRNAseq. Assuming robust sub-populations of corticotrophs exist, they should be characterised by a similar set of marker genes in all datasets. We thus sought to define a correspondence between clusters of the different datasets based on the expression of specific markers; since clustering analysis produced a different number of clusters for different studies, this correspondence could be either one-to-one or many-to-one. We determined marker genes for each dataset by identifying differentially expressed genes between all subclusters of that dataset using the Wilcoxon Rank Sum test, as implemented in the R Seurat package, considering only genes with an adjusted p-value lower than 0.05. Only genes with a log_e_ fold change greater than 0.25 and expressed in at least 20% of cells were considered for this analysis. The percentage of markers shared by different subclusters in all datasets was then compared pairwise (i.e. each cluster of dataset A vs each cluster of dataset B). In most cases, only approximately 10% (median 7.7% in datasets from males, 5.3% in datasets from females) of marker genes were found to be shared with any other cluster (Figure 2A, Supplementary Figure 2A).

**Figure 2.**
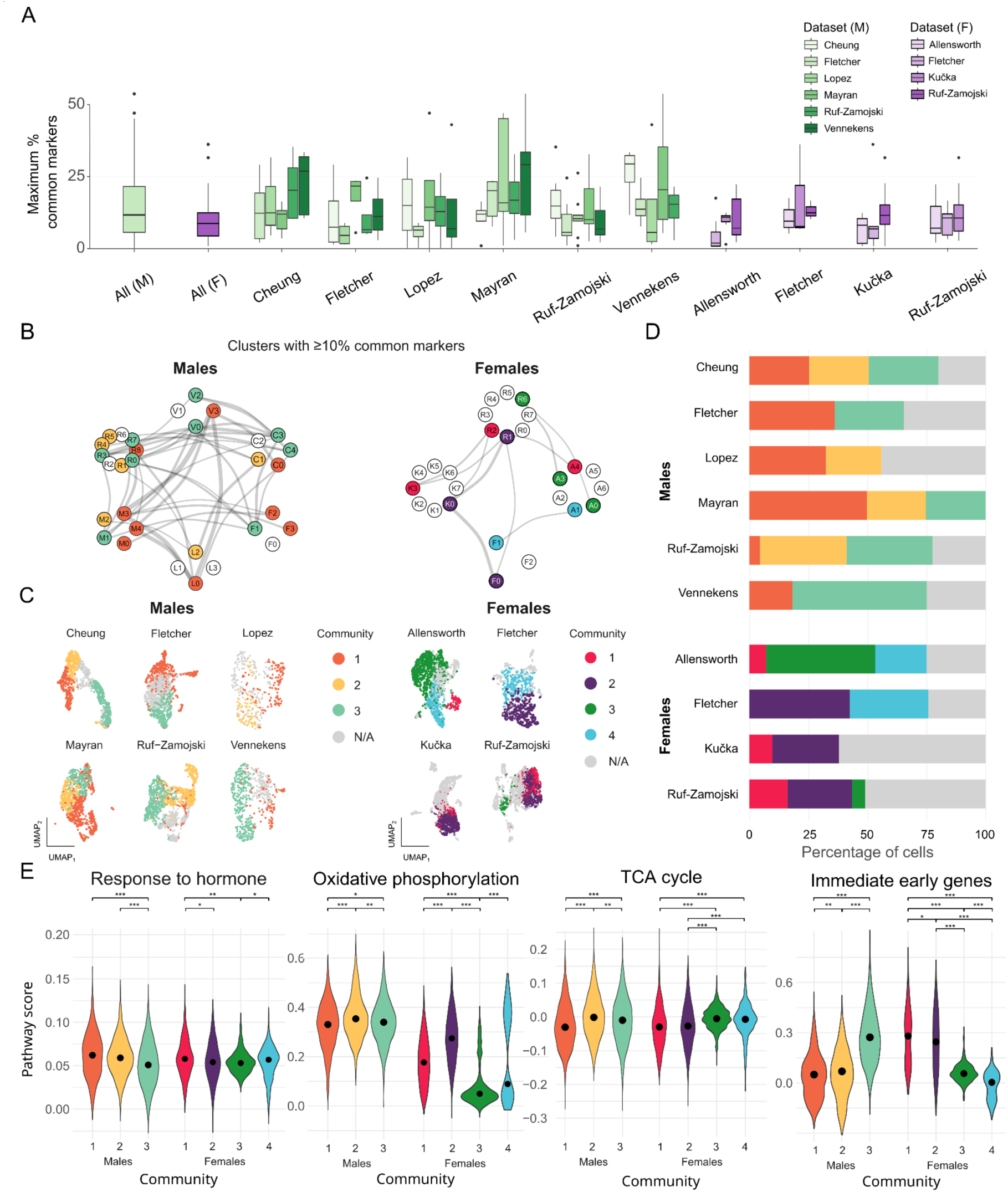
Analysis of markers of corticotrophs clusters. **A)** Maximum percentage of markers in common between all pairwise combinations of clusters for the different data sets from male animals (left, shades of green) and female animals (right, shades of purple). **B)** Graph showing the relationship between the different sub-clusters of corticotrophs. Each node represents a subcluster, indicated with the initial of the first author of the respective articles (so, for instance, cluster 2 from Fletcher 2019 is indicated as “F2”). Nodes are connected if the two clusters share at least 10% of markers; edge thickness is proportional to the % of shared markers. Nodes are coloured depending on which community they belong to, determined using the Walktrap algorithm **C)** Communities defined in B, mapped on the UMAP projections for each dataset. **D)** Percentage of cells in each community in the different datasets. E) Violin plots showing distribution of scores (circles indicate medians) for selected enriched pathways in the communities. “Response to hormones” includes genes annotated with GO:0009725 (response to hormone), GO:0019933 (cAMP-mediated signalling) or KEGG mmu04081 (hormone signalling). “TCA cycle” includes genes annotated with GO:0006099 (tricarboxylic acid cycle) or KEGG mmu00020 (citrate cycle). “Oxidative phosphorylation” includes genes annotated with GO:0006119 (oxidative phosphorylation), GO:0045333 (cellular respiration) or KEGG mmu00190 (oxidative phosphorylation). “Immediate early genes” includes genes in the list from (Tullai et al., 2007). * p<0.05, ** p<0.01, *** p < 0.001, two-tailed bootstrap p-values (males and females analysed independently).

To confirm that the marker genes effectively distinguished between clusters, we calculated pairwise Spearman’s correlation for all marker genes of each cluster, using each dataset as a reference in turn. We expected a positive correlation within clusters, meaning cells in the same cluster should show high expression of their markers, while other clusters would express these genes at lower levels. In contrast, genes marking different subclusters should show low correlation. To summarize these results, we calculated the average correlation for each cluster’s markers. Our analysis confirmed significantly higher correlations within clusters compared to between clusters (Supplementary Figure 2C, estimate (CI)_within_= 0.084 (0.072;0.095), p<0.001, mixed-effects model). However, the within-cluster correlation was significantly lower when it was calculated using markers taken from a different dataset (Figure 2D, estimate (CI)_same dataset_ = 0.034 (0.020;0.049), p<0.001, mixed-effects model). There was no effect of sex (estimate (CI)_sex_ = 0.006 (-0.023;0.037), p=0.6, mixed-effects model). Importantly, randomly shuffling the gene expression matrices resulted in a complete loss of correlation (average correlation ± SD = 0.00013 ± 0.00022, estimate (CI)_shuffle_ = -0.039 (-0.063;-0.016), p<0.001, mixed-effects model). However, when expression values were shuffled only within each cluster, correlation remained higher than between-cluster levels. This suggests that the relationship between these genes may be important for defining corticotroph subpopulations (estimate (CI)_shuffle within_ = 0.067 (0.044;0.091), p<0.001 mixed-effects model).

To visualise these relationships between the analysed datasets, we generated a graph with nodes corresponding to each of the subclusters and edges with strength proportional to the percentage of common markers; in this graph, tightly connected groups of nodes where subclusters share a higher proportion of marker genes were designated as “communities”. These communities represent sets of nodes exhibiting similar gene expression patterns, thus they can suggest potential biological similarities across datasets.

Because of the small similarity between clusters, we used a low threshold (10% marker similarity) for community detection (Figure 2B); as expected, higher percentages (15 or 20%) resulted in lower similarity and a smaller number of connections between clusters (Supplementary Figure 2B). Three communities of clusters were found in male datasets and four in the female ones (Figure 2C); importantly, each community could contain cells that we annotated in different clustering (e.g. clusters 2 and 3 from (Fletcher et al., 2019) both belong to community 1) balancing any possible over-clustering in our initial procedure. In males, the cell community 1 was found in all datasets (in orange in Figure 2C), albeit in different proportions (Figure 2D); community 3 was found in all but one dataset (in green in Figure 2C), and community 2 was found in 4 out of 6 datasets (yellow in figure 2C). In females, four communities were identified, of which two, 1 and 2, (in red and purple in Figure 2C) were common to three of the four datasets, and 3 and 4 (in light blue and green in Figure 2C) were found in two out of four datasets. The proportion of the total population belonging to each community is shown in Figure 2D; both in males and females, some clusters did not show enough similarity with any other and were thus marked as not belonging to any community (“N/A” in Figure 2D). Clusters belonging to the same community do not necessarily all share the same markers, although they do share at least 10% of markers with some other cluster in their community. Overall this low level of similarity between sub-clusters points to “local” rather than “global” similarities, i.e. within pairs of datasets rather than between all.

Overall, while the marker genes were largely dataset-specific, these results confirm their effectiveness in distinguishing corticotroph subclusters, supporting the hypothesis that these subclusters represent transient cell states rather than robust sub-populations present across all the studies.

### Cluster communities are associated to specific biological functions

As noted above, only a small number of markers were common across all clusters within each community. To further characterize these states, we opted to use the union of the marker sets, considering any gene shared by at least two clusters within the community. This approach resulted in a larger set of genes for further analysis. We then performed pathway analysis on these gene sets to gain insights into the functions of the identified communities. In males, genes overexpressed by cells in community 1 (Figure 2C, left, orange) were enriched for pathways related to the response to neuropeptides and hormone activity, specifically activating the cAMP pathway e.g. including genes such as *Pomc*, *Pou1f1*, *Gnas* and *Cga*. Other pathways enriched in these cells are related to responses to calcium, including several genes coding for proteins involved in calcium signalling, such as *Anxa1* and members of the S100 family (*S100a1*, *S100a10*, *S100a11*, and *S100a16*). Pathways associated with ribosome biogenesis and translation (KEGG:mmu03010, GO:0003735, GO:0006412) were also enriched, with multiple genes from the *Rpl* and *Rps* family, which might be important for the rapid production of signaling molecules to respond to hormonal cues.

Community 2 (Figure 2C, left, yellow) genes were associated with pathways related to cellular respiration (such as GO:0006119, GO:0042773, GO:000395), including a large number of genes belonging to the electron transport chain, such as those coding for subunits of the NADH dehydrogenase (23 genes in the *Ndufa* and *Ndufb* families), and of cytochrome c oxidase (9 genes of the *Cox* family); pathways related to translation (such as KEGG:mmu03010, GO:0003735) including 62 genes in the *Rpl* and *Rps* families, encoding for ribosomal proteins and 13 genes in the *Eif* family, encoding for elongation factors important for the initiation of translation.

Marker genes of the cells in community 3 (Figure 2C, left, green) included several members of the immediate early genes family such as *Fos*, *Fosb*, *Jun*, *Egr1* to *4*, *Ier2* and *5*, *Arc* as well as several heat-shock proteins genes (10 members of the *Hsp* family of genes and 9 of the *Dnaj* family). Notable in relation to the regulation of corticotrophs responses to hypothalamic signals is the presence of nuclear receptors such as *Nr3c1* (coding for the glucocorticoid receptor), *Nr4a1* and *Nr4a2* (immediate early genes coding for the Nur77 and Nurr1 transcription factors which are important in the regulation of *Pomc* transcription). Many of these genes are known to be activated in corticotrophs after stimulation by hypothalamic factors.

Overall, we identified three main communities that appear to represent a functional spectrum where cells in Community 1 are characterized by active hormonal signaling and responsiveness; Community 3 cells show activation of early response genes, indicating they might be in a later stage of hormonal signalling, and cells for Community 2 cells might be functioning as support or reserve.

In females, markers of Community 1 (Figure 2C, right, pink), similarly to Community 3 in males, included several immediate early genes such as *Fos*, *Fosb, Jun, Egr1, Egr2, Ier2,* heat-shock proteins (*Hsp90aa1*, *Hspa8*, *Hspb1*, *Dnaja1* and *Dnajb1)* and the *Nr3c1* gene.

Markers of Community 2 (Figure 2C, right, purple) were enriched in pathways related to response to hormone such as the receptors for hypothalamic secretagogues *Crhr1* and *Avpr1* and similarly to Community 1, a number of early response genes and nuclear receptors (*Nr3c1*, *Nr4a1* and *Nr4a2*). Genes mediating ATPase activity were also found, in particular those coding for subunits of vacuolar ATPases, which are involved in acidification of secretory vesicles (*Atp6ap1*, *Atp6v0c*, *Atp6v0d1*, *Atp6v0e2*, *Atp6v1d*).

Markers of Community 3 (Figure 2C, right, green) did not show significant enrichment for any pathway.

Finally markers of Community 4 (Figure 2C, right, blue) were limited to six genes, related to hormonal activity such as *Pomc*, the *Cpe* gene, coding for carboxypeptidase E, a protease important for POMC processing, as well as the *Chga* gene, coding for chromogranin A a protein associated with secretory granules.

In summary, similarly to what happens in males, we observed different groups of cells in females that might represent different states along a spectrum of cellular response to hypothalamic signals.

Supplementary Table 3 lists all of the enriched pathways in the different communities.

To provide a quantitative measure of functional enrichment, we calculated “module scores” for the main functional pathways described above (Figure 2E), by averaging the expression of predefined gene sets associated with each pathway, as annotated in Gene Ontology and KEGG databases, and subtracting the average expression of control gene sets with similar expression distributions. This confirmed that, despite differences in the specific genes that are expressed in the various clusters within each community, the overall functional signature of these communities has biological relevance and can be used to predict the putative functions associated with different cell states *in silico*.

### Label transfer identifies shared and dataset-specific transcriptional states

The analyses described so far are biased in considering only marker genes, a small set of genes which are, by definition, highly variable and highly expressed, or considering only pairwise relationships between genes through correlation. It is, however, much more likely that any cellular state is determined by more complex, non-linear relationships between multiple genes. To overcome these limitations, we employed an unbiased label transfer approach. We trained one multi-layer perceptron classifier for each dataset to recognise cells of that study’s clusters. Given a new cell, never seen at training time, these models can predict what cluster this cell belongs to based on the data from each study (Figure 3A)

**Figure 3.**
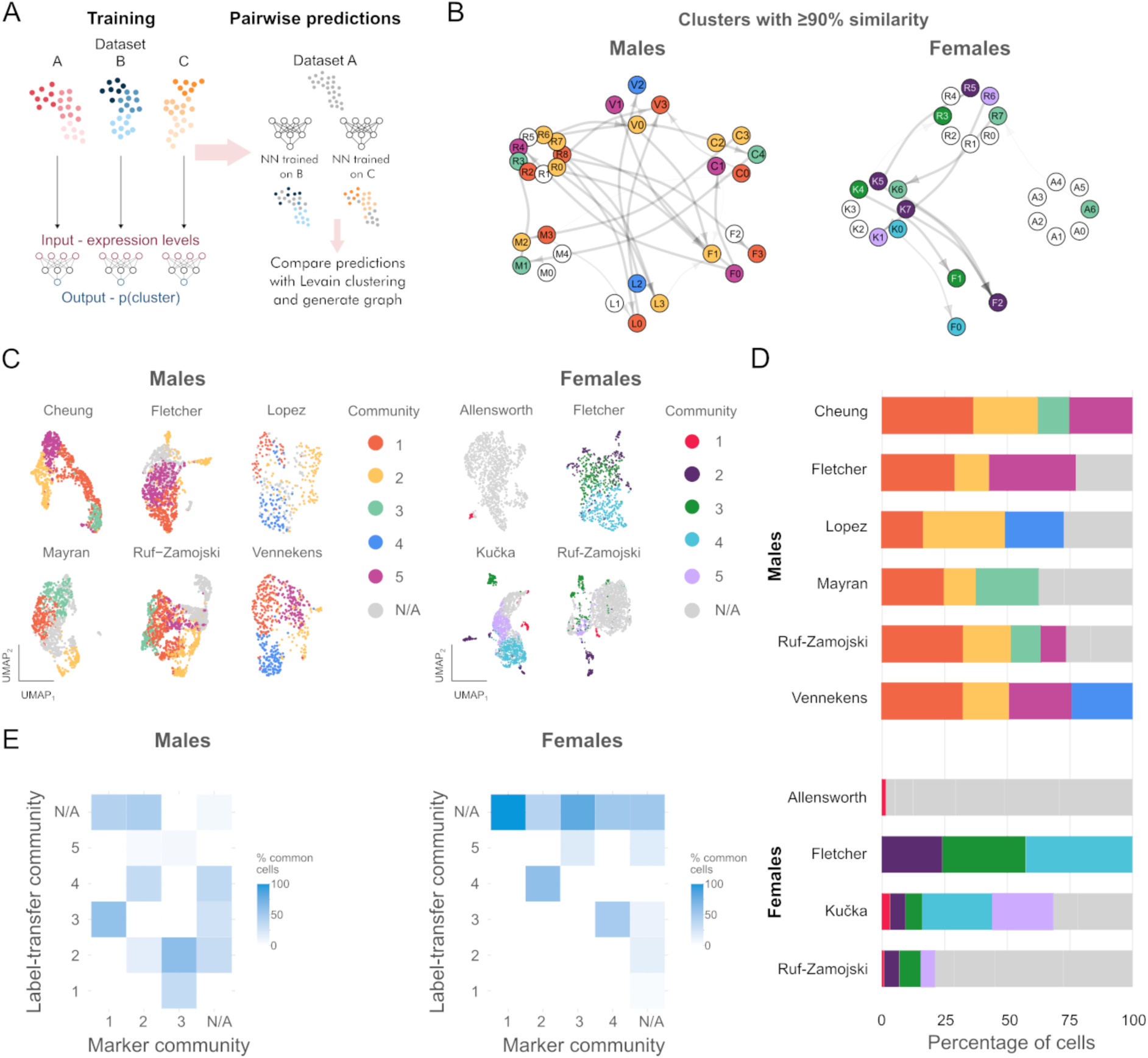
Label transfer identifies transcriptional states. **A)** Schematic of the label transfer approach. A set of neural networks, one per dataset, are trained on the expression data. Once trained, these can be used to predict labels from the other datasets. **B)** Directed graph showing the relationship between the different sub-clusters of corticotrophs. Each node represents a subcluster, indicated with the initial of the first author of the respective articles, as in Figure 2B. A connection from an input node to a receiving node indicates that at least 90% of the cells in the receiving cluster are labelled as belonging to the input cluster by the neural network trained on the input dataset; edge thickness is proportional to the % of cells. Nodes are coloured depending on which community they belong to, determined using the Walktrap algorithm. **C)** Communities defined in B, mapped on the UMAP projections for each dataset. **D)** Percentage of cells in each community in the different datasets. **E)** Confusion matrices depicting the relation between the communities defined using marker genes (see Figure 2B-D) and those defined through label transfer (Figure 3B-D).

All the models were evaluated on a test set (10% of the cells, for which the ground truth labels were known), which was held out at training time to avoid overfitting and ensure that the performance metrics are a fair reflection of the models’ generalisation ability (Figure 3A). The models performed well, with an average F1 score ± standard deviation for the test set of 0.81 ± 0.05 and an average ROC AUC of 0.80 ± 0.05 (Supplementary Table 5).

Similarly to what described above, we constructed graphs where each node corresponds to a cluster and is connected to nodes of the other datasets if more than 90% of cells have been predicted to belong to that cluster (Figure 3B; note that contrary to what seen in Figure 2B, this is a directed graph as the relationship is not necessarily symmetric); as described above, the Walktrap algorithm was used to identify cluster communities (Figure 3B-D). This identified five cluster communities in the male datasets; clusters 1 and 2 were present in all datasets, while the others were only characteristic of a subset of the datasets (Figure 3B-D). Five communities were also identified in the female datasets, although these are much less consistent between datasets. In particular, the Allensworth dataset was more dissimilar to the other three, with only a small minority of cells being mapped to other datasets. While we did not extensively investigate the reason for this, it could either be related to the low counts Overall, no community was found in all female datasets.

We then investigated the overlap between cell communities defined by the two approaches (% of common markers and label transfer, Figure 3E). In males 94% and in females 48% of the cells that were not identified as belonging to any community through markers (22% of the total cells in males and 47% in females) were assigned to a community by the label transfer process. For the cells belonging to a marker community, the label transfer assigned 48% to 67% of the cells in males and 19% to 60% in females.This suggests that both approaches capture specific transcriptional states of corticotrophs.

We explored which features were used by the classifiers to determine their predictions, using Deep Shapley additive explanations (DeepSHAP) (Lundberg and Lee, 2017), which quantify the contribution of each gene’s expression to the model’s decision. By applying DeepSHAP to our trained MLP classifiers, we identified the most influential genes for each predicted cluster and community, allowing us to compare feature importance across different models and datasets. The top 50 positively-contributing features for each cluster in each dataset are listed in Supplementary Table 6. Only less than 2% of the top 50 features that contributed to defining a cluster through the label transfer approach were also marker genes. While these top-contributing genes were not statistically enriched for any particular biological pathway, they did include genes that are known to be important for corticotroph function, such as the glucocorticoid receptor *Nr3c1*, *Pomc*, or immediate early genes such as *Fos*, *Jun* and *Egr1*. Overall, this indicates that a more complex, non-linear contributions of a large number of genes might be sufficient to describe cellular state in corticotrophs, reinforcing the idea that cells exist in a continuum of transcriptional states.

### Velocity analysis identifies conserved dynamic transitions across cell-states

To investigate whether these states can dynamically transition into one another we performed RNA velocity analysis (Bergen et al., 2020; La Manno et al., 2018). By analysing the ratio between the spliced and unspliced forms of the sequenced mRNA, vectors of transition between cell states are defined. For each dataset, we calculated RNA velocity, to determine 1) whether cells from different communities could be transitioning into each other and 2) what the direction of that transition might be (see Methods for a description of this procedure). This process was repeated both for communities defined through sharing common markers and through label transfer (Figures 4 and Supplementary Figure 3 respectively).

**Figure 4.**
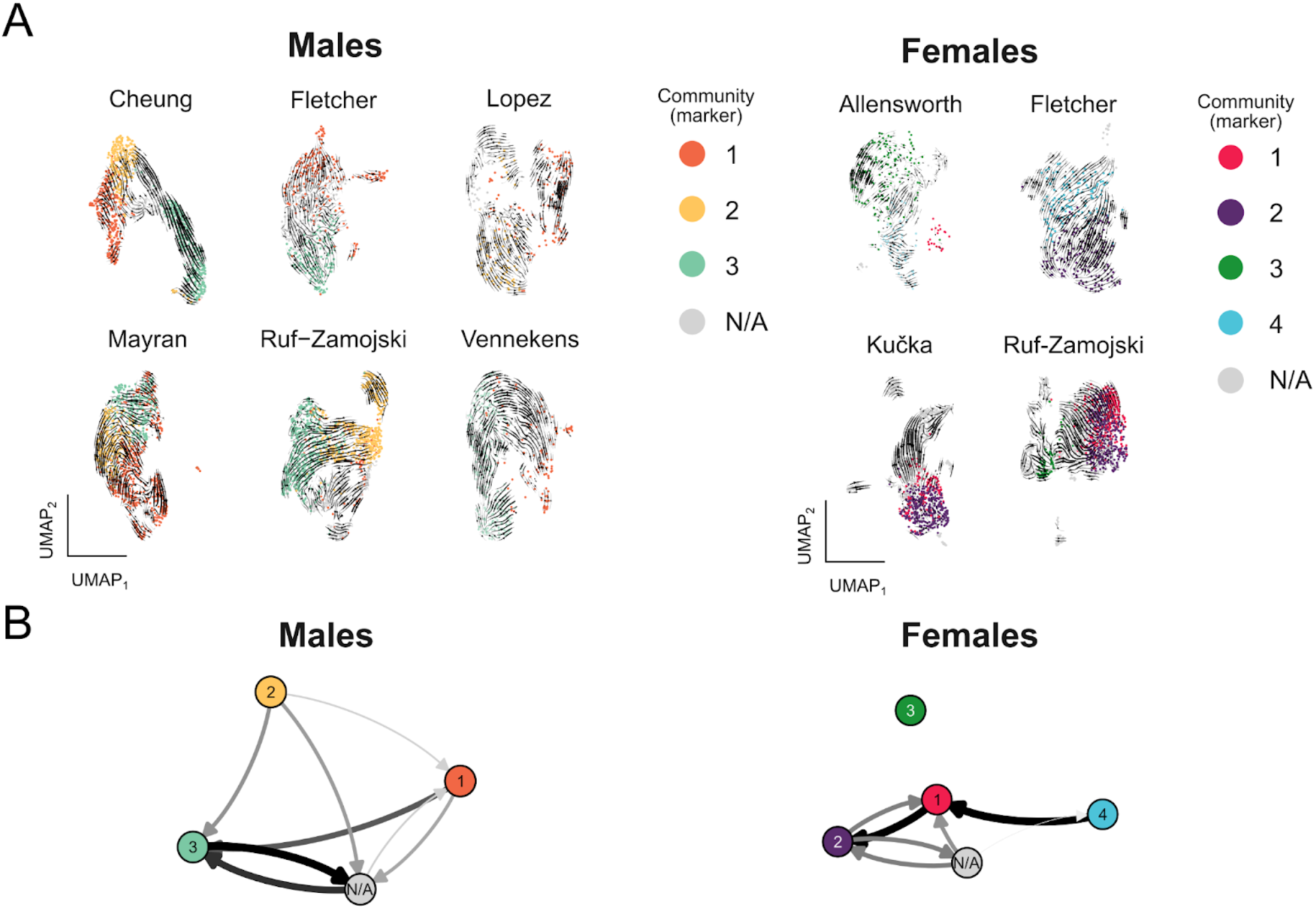
Velocity analysis identifies conserved transcriptional state transitions between studies. **A)** RNA velocity vectors superimposed over UMAP plots of the datasets; cells are coloured by cluster community, as defined by shared markes and shown in Figure 2. **B)** Directed graph showing the main direction of transitions between the different communities/cell states. Each node represents a community of clusters, coloured as in Figure 2. Connections indicate a transition between two communities, with edge thickness and darkness proportional to the probability of that transition.

In all the datasets, velocity analysis showed evident transitions between different states, including some for those cells that were not mapped to specific communities, which might indicate transition states between the ones that were identified with our analyses.

Using this information, we could generate a graph of the possible transitions for cells belonging to the communities defined either by shared marker genes (Figures 4A, 4B) or through label transfer (Supplementary Figures 3A, 3B). Despite some variability between datasets, some of these transitions were seen consistently in multiple experiments. Specifically, in males, cells from community 1, which were enriched for genes related to hormone responses transitioned towards those of community 3, enriched in early response genes, which might indicate subsequent activation of subsets of corticotrophs. A transition from community 3 to a state not belonging to a specific community was also found (n/a in Figure 4B), which could transition back to community 1, indicating corticotrophs could cycle between these two transcriptional states. Cells of community 2, which could represent a reserve state, characterised by higher expression of genes related to cellular respiration could also transition towards this unmarked state. This is mirrored in females where communities 1 and 2, expressing genes related to hormone response and early response genes, transition to each other. Community 4, expressing genes important for POMC processing and secretion also transitioned towards community 1. No specific transition was seen from or to Community 3.

Overall, both in males and in females, there was indication of similar transitions between transcriptional states relevant for the response to hypothalamic stimuli and the overall function of the HPA axis.

## Discussion

The capacity to respond to time-varying external stimuli is critical for the maintenance of body homeostasis, and the pituitary gland plays a central role in this process. This is particularly significant in the context of hormonal stress responses, where the body needs to rapidly respond to unpredictably changing external stimuli of different strength, length and type. Corticotrophs of the anterior pituitary are the cells responsible for the secretion of adrenocorticotrophic hormone (ACTH), the main controller of adrenal cortisol production and secretion, and they have been shown to display heterogeneous functional behaviour (Romanò et al., 2017; Liang et al., 2011; Takigami et al., 2008; Lugo and Pintar, 1996; Corcuff et al., 1993; Childs and Burke, 1987).

Heterogeneity at the single-cell level is a ubiquitous property of biological systems. While some of this variability arises from stochastic cellular processes, it is increasingly recognised as functionally relevant rather than simply an artifact of transcriptional noise (Wada et al., 2021). In fact, heterogeneity can confer advantages at the population level by promoting emergent behaviors that enable collective cellular functions beyond the capabilities of individual cells (Paszek et al., 2010). For example, heterogeneity is at the base of bet-hedging strategies in bacterial populations, enabling a subset of cells to enter a dormant, antibiotic-tolerant state when exposed to an antibiotic; once the drug is removed, these cells allow the regrowth of the bacterial population, despite not having acquired any genetic resistance to antibiotics (Balaban et al., 2004). In the vascular system, heterogeneity can increase the dynamic range of responses of the cell population because small groups of endothelial cells are sensitive to restricted ranges of acetylcholine concentrations, allowing the system to respond to a much larger range of stimuli without being saturated by random fluctuation in basal acetylcholine release (Wilson et al., 2016). Similarly, mouse fibroblasts respond to tumour-necrosis factor α (TNFα) in a digital-analogue manner, with different patterns of TNFα stimulation resulting not only in responses of different intensity, but also in the recruitment of a different proportion of the cell population (Adamson et al., 2016); this behaviour is similar to what we previously reported in corticotrophs in response to CRH and AVP (Romanò et al., 2017) that could allow the population to respond to a more extensive, diverse range of stressors. Our functional data showed that the heterogeneous behaviour seen at the population level, with cells showing a range of different responses to the same stimulus, was not mirrored by single cells, which instead responded deterministically to repeated challenges with the same stimulus in the time-frame of our experiments. This led us to hypothesize that there might be different sub-types of corticotrophs, which might be defined by specific transcriptional signatures.

To test this hypothesis, here we have undertaken a comprehensive analysis of previously published scRNAseq datasets from rodent pituitary with the aim of defining whether stable transcriptional sub-populations of corticotrophs exist, or if they exist in dynamic transient states. This aims to provide insights into whether these subpopulations could account for the heterogeneous functional behaviours observed in these cells. Our rationale was that should stable subpopulations exist, they should be detectable in any of the datasets we analysed, in comparable proportions. However, our analysis revealed a more intricate scenario, where corticotrophs exist in several transcriptomic states and are able to transition between them. Thus, when looking at the temporal snapshots obtained from scRNAseq we see a study-specific pattern of transcriptomic states; this means we are able to define some correlation between different studies, when cells happen to be in a similar state at that specific time of the experiment, as well as detecting their trajectory of transition

Our initial hypothesis was strongly influenced by our previous work on calcium responses to secretagogues in corticotrophs; there, we showed that while different cells responded heterogeneously to the same stimulus, single-cell responses were deterministic and remained constant during the time frame of our experiments (Romanò et al., 2017). However, while those experiments were dynamic, they were, for technical reasons, generally limited to less than three hours; we might thus have been observing short stable states of these cells. Indeed, work on other pituitary cell types has shown complex long-term changes in transcription (Featherstone et al., 2016) and linked transcriptional patterns of hormonal genes to calcium responses (Harper et al., 2020; Villalobos et al., 2002) and thus, at least indirectly, to hormonal secretion. Furthermore, these experiments demonstrated that cells could transition between transcriptional states; for example, rat pituitary GH3 cells can be sorted into “High” and “Low” populations based on *Prl* gene transcription; when assayed the following day, the “Low” population has reverted to a mixture of “High” and “Low” cells, via calcium-dependent modulation of histone acetylation, possibly dependent on the state of surrounding cells (Harper et al., 2020). Similar to what we previously observed in primary cells (Romanò et al., 2017), this might serve as a reserve mechanism, allowing non-active non-secreting cells to be engaged in responses to unpredicted stimuli, while active secreting cells can provide a basal level of hormone production.

Our analysis compared transcriptional signatures between datasets using a marker gene-based strategy and a whole transcriptome label transfer approach. In the first case, after sub-clustering the cells using the commonly used Louvain clustering method, we performed all pairwise comparisons between the marker genes of each cluster in each dataset. This revealed a surprisingly low similarity, with often less than 10% of markers in common between clusters of different datasets. A similar observation has been made when comparing the markers reported by scRNAseq studies on pancreatic β-cells, where similarity in marker genes of heterogeneity was extremely low (Mawla and Huising, 2019). There are several possible explanations for this low similarity; just like for the insulin gene in the case of β-cells, the transcriptome of corticotrophs is heavily skewed towards the transcription of *Pomc*. Indeed, *Pomc* represented between 3 and 25% of the corticotroph transcriptome in the datasets we considered, therefore possibly masking differences in lowly-expressed genes. Furthermore, scRNA-seq data is highly sparse; on average, 86% and in one case, 97% of the counts in the analysed datasets were 0, leading to issues for the statistical analysis of these datasets, the so-called “dropout problem” (Church et al., 2022; Kharchenko et al., 2014). This, however, could also be a result of classifying subpopulations using a biased selection of “marker genes”, which are, by definition, highly variable and highly expressed. To extend our analysis to the complete gene signature of each cell, we further employed a label transfer strategy (Fidanza et al., 2020; Stumpf et al., 2020) by creating a series of neural network classifiers which can map clusters from one study to another. This resulted in a mapping which only partially corresponded to what we found with the marker-based analysis. Contrary to what happened when a similar approach was applied to transfer labels for different cell types (Fidanza et al., 2020), the vast majority of the genes used by the neural networks to classify clusters were not markers. This suggests that the overall transcriptional fingerprint of the cell can hold important information to define cell states, which can be lost when only analysing highly expressed-highly variable genes. The fact that the two methods identified only partially overlapping communities of markers, supports the idea that the limit between these states is somehow arbitrary, and these are more accurately represented as a continuum of transcriptional states rather than discrete, rigidly defined cell subtypes or states.

The sparsity of scRNAseq datasets means that several genes which have been experimentally validated as key components of corticotrophs and are thus biologically relevant markers, such as several ion channels, are poorly detected in these experiments, due to their low level of expression. For example, a large body of literature shows the important role of the large conductance calcium-activated potassium channels BK in corticotrophs and other pituitary cell types (Duncan et al., 2015; Tabak et al., 2011; Miranda et al., 2003). However, *Kncma1*, which encodes for the alpha subunit of the channel, is not found in an average 91.5% of corticotrophs in the datasets we analysed, and is detected with a count of 1 in 7.4%, with a maximum count of 7 in only two cells in all the datasets. Given that very small changes in the number of open BK channels at the cell membrane are predicted to have a significant physiological effect on corticotroph physiology (Duncan et al., 2022), this highlights a critical limitation of scRNA-seq for identifying functionally important genes. This is a particular problem for genes with moderate to low expression levels that can still exert significant physiological effects, an issue that becomes even more evident when trying to distinguish cell sub-populations. Despite these issues, correlation analysis confirmed a non-random association of the marker genes in the clusters, and the transcriptomes of groups of clusters that shared common markers were more correlated than those that did not, indicating this was not an intrinsic issue with how these marker genes were discovered.

Pathway analysis based on common marker genes revealed both in males and females the presence of cells that are in a transcriptional state characterised by genes, such as those involved in the cAMP/PKA pathway or in calcium signalling, involved in the very immediate intracellular response to hypothalamic secretagogues. Other cells show a strong signature of early response genes, possibly indicating a subsequent stage of signalling. Finally, a group of cells that show a different signature, characterised by genes involved in cellular respiration, which might indicate a “reserve” state. Indeed, our recent bulk transcriptomics analysis of corticotrophs shows that similar pathways are central for the functional changes happening during chronic stress and recovery (Duncan et al., 2024). Dynamic transition across cell-states, calculated using Velocity analysis suggests that corticotrophs might be transitioning between transcriptional states in a way which would allow the system to quickly recruit cells to respond to unpredictable stimuli. These transitions are observable both in males and females, in mice as well as rats, indicating they are a fundamental property of how the corticotroph population works. This suggests that different groups of corticotrophs might be active at different times, possibly to ensure robustness in the system, similarly to what has been recently shown in the case of the GnRH neurons controlling fertility (Yeo et al., 2025). Multiple factors tied to the specific status of the HPA axis at the time of the experiment might be driving these transitions; for example, the complex circadian and ultradian dynamics of this system might be influencing the proportion of cells in different states. A similar dynamic picture is seen when considering communities of clusters identified through transfer learning, despite differences in the exact membership of each cell. This is likely a result of the fact that clustering is an ill-posed problem, and the number of clusters resulting from any clustering algorithm is arbitrary (Jain, 2010) and strongly dependent on the features that are used. While we do acknowledge that there is no “correct” set of clusters, our comparison should still be valid, as the general relationships between the clusters should be consistent across datasets if relevant biological signals are present.

Overall, our analysis supports the idea of dynamically changing secretory cells, similarly to what has been reported in rat lactotrophs, where a large proportion of cells secrete prolactin in an intermittent manner, with the amount secreted by the same cell varying over the course of several days, and correlated to fluctuations in hormonal transcription (Castaño et al., 1994, 1996). This basal variability in secretion can also be a mechanism at the base of the recruitment of different subpopulation of cells with varying levels or patterns of stimulus as we and other showed in corticotrophs (Romanò et al., 2017; Takigami et al., 2008) and in other cell types (Benninger and Kravets, 2022; Adamson et al., 2016; Salomon and Meda, 1986).

Finally, this study also contributes to highlighting important issues linked to the inherently noisy and low-sensitivity nature of current scRNAseq technologies. Each study only shows a single snapshot in time of a complex, strongly time-varying process so it is important to investigate multiple independent replicates. Furthermore, better reporting of analysis methodologies and choice of parameters will ensure workflows that are fully reproducible, and lead to generalisable conclusions. Indeed, several studies have identified transcriptional sub-types of pituitary cells (Sheridan et al., 2024; Cheung et al., 2018, 2023; Zhang et al., 2020, 2022; Ruf-Zamojski et al., 2021) often based on single experimental units. Pseudoreplication is a major issue in scRNAseq experiments, which is exacerbated by the large batch to batch variation in these experiments (Squair et al., 2021; Zimmerman et al., 2021), increasing the risk that some of the observed cell subpopulations might be simply an artefact of underpowered studies. The requirements for number of replicates in bulk RNA sequencing has been previously explored (Schurch et al., 2016), but similar systematic studies for scRNAseq are currently missing. Furthermore, a large number of largely arbitrary parameter choices need to be performed when analysing these datasets, many of which are either not reported or left to the software default; caution should therefore be taken when comparing data analysed using different pipelines even when the same software packages have been used since different versions might produce inconsistent results (Rich et al., 2024; Soneson and Robinson, 2018). This is well exemplified by the number of pituitary cells that we obtained in our analysis after QC filtering; while we could generally obtain a number of cells equal or at least very close to what published for each dataset when using the QC methods reported in the respective publications, we could not find a unified single set of criteria that allowed us to obtain the reported number of cells for all of the datasets.

Overall, our analysis provides compelling evidence that, in their basal state, corticotrophs do not exist in fixed sub-populations but rather transition dynamically between a gradient of transcriptional states characterised by the expression of biologically relevant sets of genes, a feature that could enhance the ability of this cell population for context-dependent response to stressors. A key challenge for the future will be to unravel how these transitions between transcriptional states translate into specific, and functionally significant, differences in cellular responses. This will necessitate the development of novel experimental paradigms that allow dynamic, real-time and simultaneous measurements of hormone secretion, signaling dynamics, and other relevant physiological phenomena. Furthermore, elucidating the regulatory mechanisms governing these transitions, including the roles of epigenetic modifications, signaling pathway crosstalk, and the influence of the cellular microenvironment, will be crucial for a comprehensive understanding of corticotroph plasticity and its contribution to the fine-tuned regulation of the HPA axis in health and disease.

## Funding and Acknowledgments

This study was supported by Medical Research Council (MRC) grant MR/V012290/1 to NR, PJD, PLT and MJS. ZY is supported by a studentship funded by Zhejiang University - University of Edinburgh Institute - Biomedical Sciences. LL is supported by an Edinburgh Doctoral College Scholarship from the Deanery of Biomedical Sciences at the University of Edinburgh. ES was supported by funding from the Biomedical Teaching Organisation at the University of Edinburgh. The authors would like to thank Dr Antonella Fidanza for critical discussion and proofreading of the manuscript.

