## Supplementary Figures for "Dynamic transcriptional heterogeneity in pituitary corticotrophs"

### Dynamic transcriptional heterogeneity in pituitary corticotrophs - Supplementary figures

Supplementary Figure 1

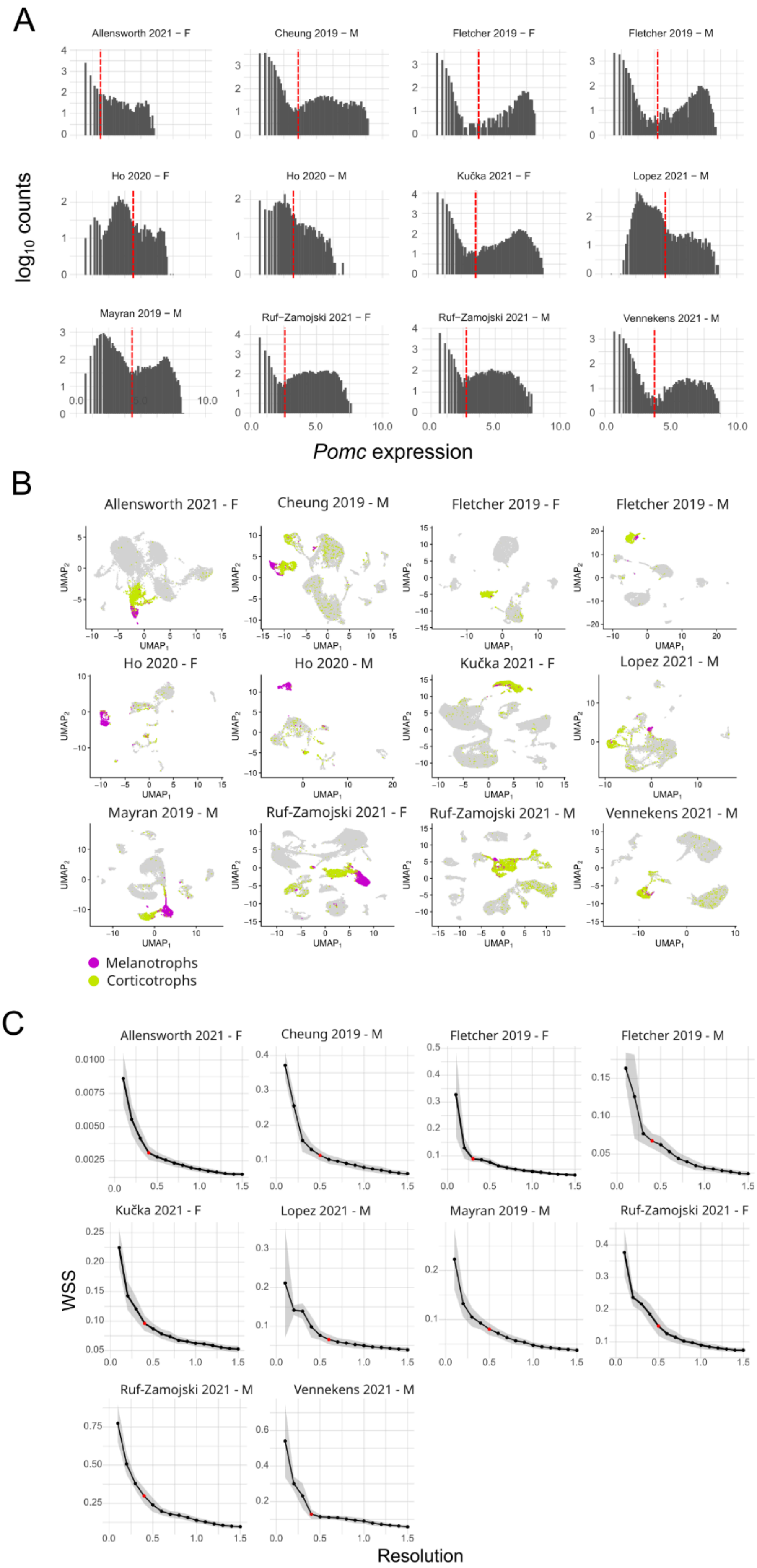

**Supplementary Figure 1** - **A)** Histogram of the levels of *Pomc* gene expression. The red line shows Otsu's threshold used for dividing the cells into *Pomc*$^{+}$ and *Pomc*$^{-}$. The same process was used for *Pax7* and *Pcsk2*. **B)** UMAP projections of the datasets, highlighting *Pomc*$^{+}$/*Pax7*$^{-}$/*Pcsk2*$^{-}$ (corticotrophs, green) and *Pomc*$^{+}$/*Pax7*$^{+}$/*Pcsk2*$^{+}$ cells (melanotrophs, purple). **C)** Plot of the average within-cluster sum of squares (WSS) depending on Louvain's clustering resolution. Grey bands show WSS mean ± SD from 100 clustering trials (see Methods for further details).

Supplementary Figure 2

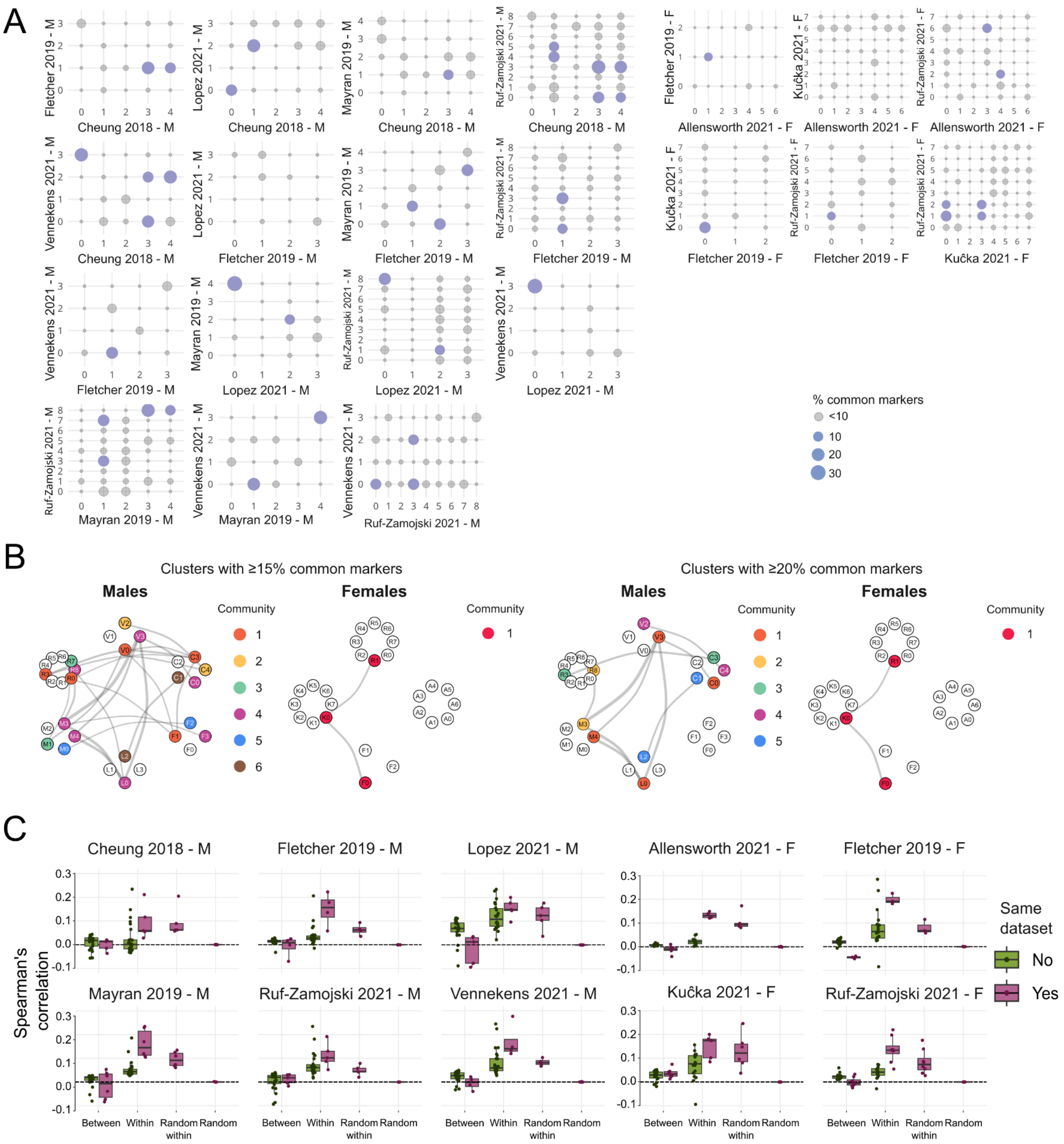

**Supplementary Figure 2 - A)** Dot plot showing the percentage of markers in common between all combination of clusters between pairs of datasets. The size of the dots is proportional to the percentage of markers; grey dots show less than 10% markers in common. **B)** Community graphs, as in Figure 2B, but for higher thresholds of common markers (left, 15%, right 20%). **C)** Boxplot summarising the results of the correlation analysis. Each subpanel represents a study and boxplots show the average Spearman's correlation in the expression of the markers from the same cluster ("Within", second set of bars) or from different clusters ("Between", leftmost two bars) in data coming from either the same (purple bars) or different datasets (green bars). Rightmost groups show within cluster correlation when data are randomly shuffled within each cluster ("Random within") or randomly shuffled globally ("Random"). Data points are overlaid on the boxplots, midline shows median, whiskers show inter-quartile range.

Supplementary Figure 3

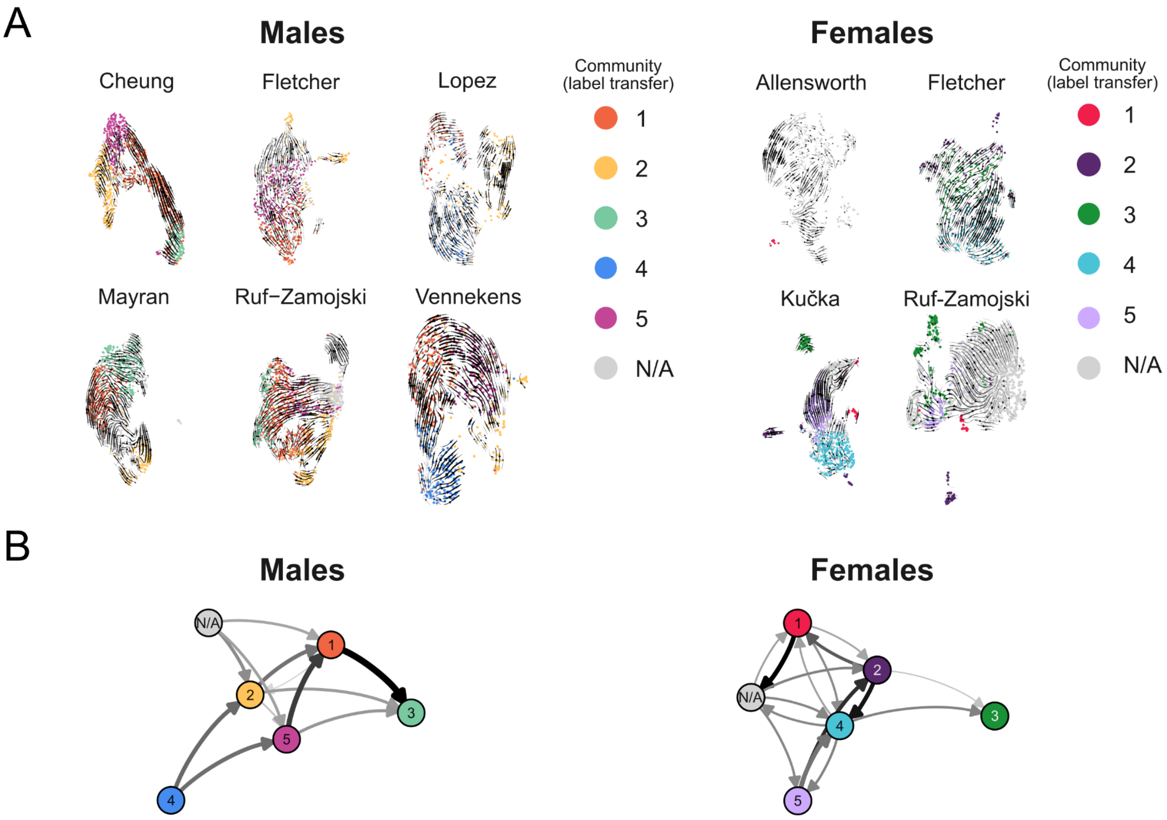


**Supplementary Figure 3 - A)** RNA velocity vectors superimposed over UMAP plots of the datasets; cells are coloured by the communities defined by label transfer, as shown in Figure 3. **B)** Directed graph showing the main direction of transitions between the different communities/cell states. Each node represents a community of clusters, coloured as in Figure 3. Connections indicate a transition between two communities, with edge thickness and darkness proportional to the probability of that transition.
