## Supplementary Tables Legends for "Dynamic transcriptional heterogeneity in pituitary corticotrophs"

### Dynamic transcriptional heterogeneity in pituitary corticotrophs - Supplementary Tables legends

**Supplementary Table 1 -** Metadata for the datasets used in this study, including source for data and sample description.

**Supplementary Table 2 -** Quality control limits reported in the datasets. Parameters that were not reported in the studies are indicated with N/A.

**Supplementary Table 3 -** Enriched pathways from KEGG, GO:Molecular Function (GO:MF) and GO:Biological Pathway (GO:BP) annotations in each of the cluster communities defined by clusters sharing more than 10% markers.

**Supplementary Table 4 -** Tuned hyperparameters for the ANNs used for label transfer.

**Supplementary Table 5 -** Evaluation of the ANNs (F1 score and ROC AUC) on the held-out test set.

**Supplementary Table 6 -** Top 50 DeepSHAP features describing the cluster communities defined through label transfer. Genes that are also markers for the specific clusters are indicated by an asterisk after their name.
